# Inhibitory cell populations depend on age, sex, and prior experience across a neural network for Critical Period learning

**DOI:** 10.1101/650051

**Authors:** Joseph V Gogola, Elisa O Gores, Sarah E London

## Abstract

In many ways, the complement of cell subtypes determines the information processing that a local brain circuit can perform. For example, the balance of excitatory and inhibitory (E/I) signaling within a brain region contributes to response magnitude and specificity in ways that influence the effectiveness of information processing. An extreme example of response changes to sensory information occur across Critical Periods (CPs). In primary mammalian visual cortex, GAD65 and parvalbumin inhibitory cell types in particular control experience-dependent responses during a CP. Here, we test how the density of GAD65- and parvalbumin-expressing cells may inform on a CP for complex behavioral learning. Juvenile male zebra finch songbirds (females cannot sing) learn to sing through coordinated sensory, sensorimotor, and motor learning processes distributed throughout a well-defined neural network. There is a CP is for sensory learning, the stage during which a young male forms a memory of his “tutor’s” song, which is then used to guide the young bird’s emerging song structure. We quantified the effect of sex and experience with a tutor on the cell densities of GAD65- and parvalbumin-expressing cells across major nodes of the song network, using ages that span the CP for tutor song memorization. As a resource, we also include whole-brain mapping data for both genes. Results indicate that inhibitory cell populations differ across sex, age, and experiential conditions, but not always in the ways we predicted.

## Introduction

Balanced excitatory and inhibitory (E/I) signaling is a widespread neural feature that facilitates informational integration and cognitive function (Zhou and Yu, 2018). E/I balance typically stabilizes across development to create efficient processing networks with controlled neural plasticity (Rubenstein and Merzenich, 2003; Wehr and Zador, 2003; Isaacson and Scanziani, 2011). In sensory brain areas, E/I balance is particularly important for increasing the selectivity and responsive properties to specific stimuli (Moore and Wehr, 2013; Natan et al., 2017; Wood et al., 2017; Vickers et al., 2018). An extreme example of shifts in response to sensory stimuli occur across Critical Periods (CPs), restricted developmental phases when a specific experience has profound and lasting effects on a particular brain system and patterns of resulting behavior (Knudsen, 2004). E/I balance may therefore play an especially essential role in neural plasticity across CPs.

Like all songbirds, zebra finches (*Taeniopygia guttata*) learn their song. In this species, only males can sing, females never sing (Zann, 1996). Despite hearing song all day every day, male zebra finches can only use song they hear during the juvenile phase that spans approximately Posthatch day (P) 30-65 to shape the structure of their own song (**Fig 1A**; Eales, 1985, 1987; Slater et al., 1988; Böhner, 1990; Roper and Zann, 2006). This limited learning phase meets the criteria for a CP because males that do not hear song P30-65 can use a song they experience later to guide their song structure; learning song prevents future learning whereas lack of learning permits late learning (Morrison and Nottebohm, 1993; London, 2017). This indicates that song is processed differentially based on sex, age, and prior tutor experience.

**Figure 1.**
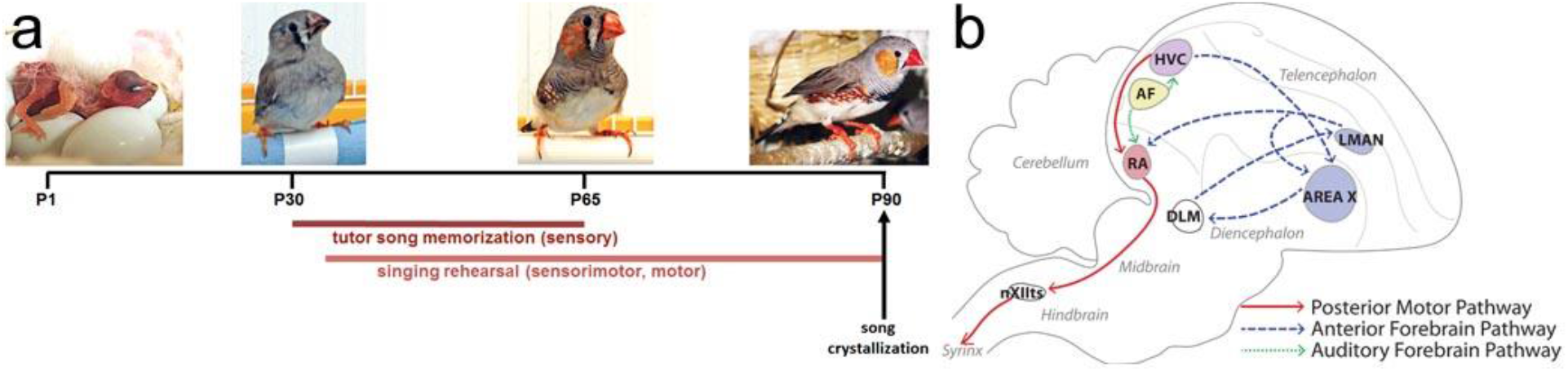
Zebra finch system for developmental song learning. **a)** Timeline of Posthatch (P) development with male images at ages relevant to tutor song memorization, and showing the multiple types of learning required for tutor song copying and adult song crystallization. **b)** Simplified schematic of the brain network for song learning and production. Major brain areas of the network quantified here are shaded. Blue = nodes necessary for sensorimotor learning during development, Area X and the Lateral Magnocellular nucleus of the Anterior Nidopallium (LMAN). Red = motor output pathway, including the Robust nucleus of the Arcopallium (RA). Purple = node in both sensorimotor and motor pathways (HVC, proper name), Yellow = Auditory forebrain (AF) required for tutor song memorization, which has three major components (not shown): primary auditory cortex Field L, and higher order association areas caudomedial nidopallium (NCM) and caudal mesopallium (CM).

The entirety of the developmental song learning process requires integrated and distributed learning across a network of brain regions (**Fig 1B**). The song network can be roughly divided into functionality for three types of learning (Brainard, 2004; Deregnaucourt et al., 2004; Fee et al., 2004; Theunissen et al., 2004; Williams, 2004; Hahnloser and Kotowicz, 2010; London, 2017). The foundation of learned song structure is the memory a young male forms of an adult “tutor” male’s song, which normally occurs P30-65 and which requires the auditory forebrain (composed of primary auditory cortex Field L, and higher order association areas caudomedial nidopallium (NCM) and caudal mesopallium (CM; Eales, 1985, 1987; Roper and Zann, 2006; London and Clayton, 2008; Ahmadiantehrani and London, 2017). A process of sensorimotor error correction then incorporates the tutor song memory into the bird’s immature vocalizations via the Anterior Forebrain Pathway comprised of HVC (proper name), Area X, and the lateral magnocellular nucleus of the nidopallium (LMAN). Finally, through intense motor rehearsal, that song becomes highly stereotyped, or crystallized, such that each male sings one song his entire adult life, using the Posterior Motor Pathway of HVC and the robust nucleus of the arcopallium (RA; Simpson and Vicario, 1990)).

Neuroanatomical, epigenetic, molecular and physiological response properties emerge and become selective differently within the different nodes of this network as developmental song learning proceeds, and a functional contribution of local inhibitory networks has already been described in the sensorimotor song region HVC (proper name; Vallentin et al., 2016). It is therefore possible that they establish and maintain E/I balance across the network.

There are multiple inhibitory cell subtypes in the brain, and diversity within each of them (Murphey et al., 2014; Blankvoort et al., 2018). Here, we focused on subtypes defined by GAD65 (a GABA-synthesizing enzyme) and parvalbumin (a calcium-binding protein) expression because of their central roles in controlling the CP for plasticity in the mammalian primary visual cortex, the best-studied CP (Hensch et al., 1998; Takesian and Hensch, 2013; Davis et al., 2015). We asked how GAD65- and parvalbumin-expressing cell populations changed based on age, sex, and prior tutor experience across major nodes of the network for developmental song learning. The experimental design was guided by evidence that a CP for developmental song learning controls the ability to memorize tutor song, and that tutor song memorization requires higher order association areas of the auditory forebrain (Morrison and Nottebohm, 1993; London and Clayton, 2008; Mori and Wada, 2015; Yanagihara and Yazaki-Sugiyama, 2016; Ahmadiantehrani and London, 2017; London, 2017). We profiled brain-wide expression patterns for all of the experimental groups, and quantified cell density within major nodes of the song network. Our results support the broad hypothesis that inhibitory cell populations may contribute to the emergence of song processing specificity across the song network, though in sometimes unexpected ways. Future work can expand on these first findings by investigating additional inhibitory cell subtypes and measuring dynamic synaptic function of these cell populations.

## Methods

All procedures were conducted in accordance with the National Institute of Health guidelines for the care and use of animals for experimentation and were approved by the University of Chicago Institutional Animal Care and Use Committee (ACUP #72220).

### Experimental animals

All birds used in this study were hatched in in-house breeding aviaries where males and females of all ages were housed on a 14 h:10 h light:dark cycle, with seed and water provided *ad libitum*. Birds reared in the ‘Normal’ condition were allowed to remain in their home aviary until collection at P25, P45, or P65. Birds that were raised for the ‘Isolate’ condition were removed from their home aviaries the day after they fledged, P21-P23, and lived with two adult females in a sound attenuation chamber until collection at P25, P45, or P65. The Isolate condition prevents juveniles from hearing song (female cannot sing), but they are exposed to conspecific vocalizations in the form of female calls and experience social interactions. We assayed males and females for all three ages and both conditions, n=3 for each combination of age, sex and tutor experience condition.

### In situ hybridization

Whole brains were rapidly dissected, embedded in OCT (Fisher Healthcare), and flash frozen on dry ice. Brains were stored at −80°C until sectioning into 12μm sagittal slices on a cryostat in a series so that adjacent slides from each brain could be processed for GAD65 and parvalbumin across samples. The same series was used consistently across subjects for each probe.

We hybridized with antisense-configured riboprobes generated from two zebra finch ESTs (GenBank accession numbers CK310366 and FE713884, for GAD65 and parvalbumin, respectively). Briefly, EST plasmids were grown in Luria Bertani (LB) media overnight at 37°C and purified before further use (QIAprep Spin Miniprep Kit, Qiagen). Plasmids were linearized with PauI (New England Biolabs) restriction enzyme at 37°C, then used as templates for *in vitro* transcription with RNA T3 polymerase and DIG-labeled NTPs (Roche) to generate DIG-labeled riboprobes. Probes were purified before use (RNeasy Mini Kit, Qiagen), and concentration was assessed via dot blot (Roche).

Processing followed prior protocols (London et al., 2003; London et al., 2009). Sections were first dried at room temperature and then fixed for 10 min in 4% paraformaldehyde (pH 7.4). They were rinsed three times in 0.025M KPBS (pH 7.4), equilibrated in 0.1 M Triethanolamine (TEA) for 3 min, then treated with 0.25% v/v acetic anhydride in TEA for 10 min. Sections were washed twice in 2X SSC and dehydrated in a graded ethanol series. After drying at room temperature, sections were hybridized for 16 hr at 65°C in hybridization solution (50% formamide, 2X SSPE [pH 7.4], 2 mg/mL tRNA, 1 mg/mL bovine serum albumin, 300 ng/mL polyadenylic acid, 0.1M DTT) containing 400ng of either GAD65 or parvalbumin antisense-configured riboprobes. Following hybridization, sections were rinsed in 2X SSC at room temperature to remove coverslips, and then high-stringency washed in 0.1X SSC/50% formamide once and 0.1X SSC twice, all at 65°C. Sections were blocked for 1 hr at room temperature (Roche), then rinsed three times in GBA (100mM Tris pH 7.5, 165 mM NaCl) and incubated 2 hr at room temperature with a 1:5000 dilution of alkaline phosphatase-conjugated anti-DIG primary antibody in block solution (Roche, cat #11-082-736-103, cat #11-585-762-001). Sections were then rinsed four times in GBA and once in GBB (100mM Tris pH 9.5, 100mM NaCl, 100mM MgCl_2_) before incubating with BCIP/NBT alkaline phosphatase detection substrate (Sigma, cat #B5655), and rinsing in deionized water before cover-slipping in aqueous mounting media.

### Imaging and quantification

All images were captured using microscopes at the University of Chicago Integrated Light Microscopy Core Facility. Images for each region that was quantitated (Field L, NCM, CM, HVC, Area X, LMAN, RA) were obtained using a 4X objective on an Olympus IX81 microscope (Olympus Corporation of the Americas, Center Valley, PA) with a Hamamatsu Orca Flash 4.0 sCMOS camera (Hamamatsu Photonics, Skokie, IL) running Slidebook 5.0 software (Intelligent Imaging Innovations). Images were captured to include the song region of interest and surrounding brain region, used as background control. All images were captured and analyzed at the same magnification.

For each and all images, a threshold was applied in FIJI (National Institutes of Health) to exclude background staining. We then used the particle analysis function to obtain data only for positively-stained cells in each region. Quantification was done within song areas and adjacent surrounding brain regions to control for inter-section variation in staining intensity as follows: auditory forebrain (Field L, NCM, and CM) and Hippocampus (HP); HVC and nidopallium; RA and arcopallium; LMAN and nidopallium; Area X and medial striatum. LMAN could not be consistently differentiated between core and shell components across sections, so the entire region was quantified as one unit. We did not quantify Area X for females as they lack an easily identifiable Area X-like structure within the medial striatum with this staining (but see London et al., 2003). Within the auditory forebrain of each bird, we separately analyzed the one primary thalamorecipient region (Field L2a, hereafter referred to as “Field L”) and two secondary auditory regions (caudomedial nidopallium, NCM; caudal mesopallium, CM), only from midline to 1mm lateral of midline. Neuroanatomical landmarks and specific boundaries used for consistent quantification of these regions were aided by the Zebra Finch Expression Brain Atlas (ZEBrA, Oregon Health and Science University; zebrafinchatlas.org).

Data were first calculated as cell counts within the song area divided by the quantified total area to give a cell density measure. Each cell density measure of a region was then divided by a cell density measure of the adjacent non-song region, to give us a normalized cell density measure for each section. Normalization via adjacent regions was done in order to control for potential variations in tissue background or staining efficiency. These normalized section measures were averaged by song area within each subject to give one normalized average cell density measure per subject per song network node (Mello et al., 1992; Kelly et al., 2018). The normalized average cell density is the measure used in the following statistical analysis and in **Fig 2**.

**Figure 2.**
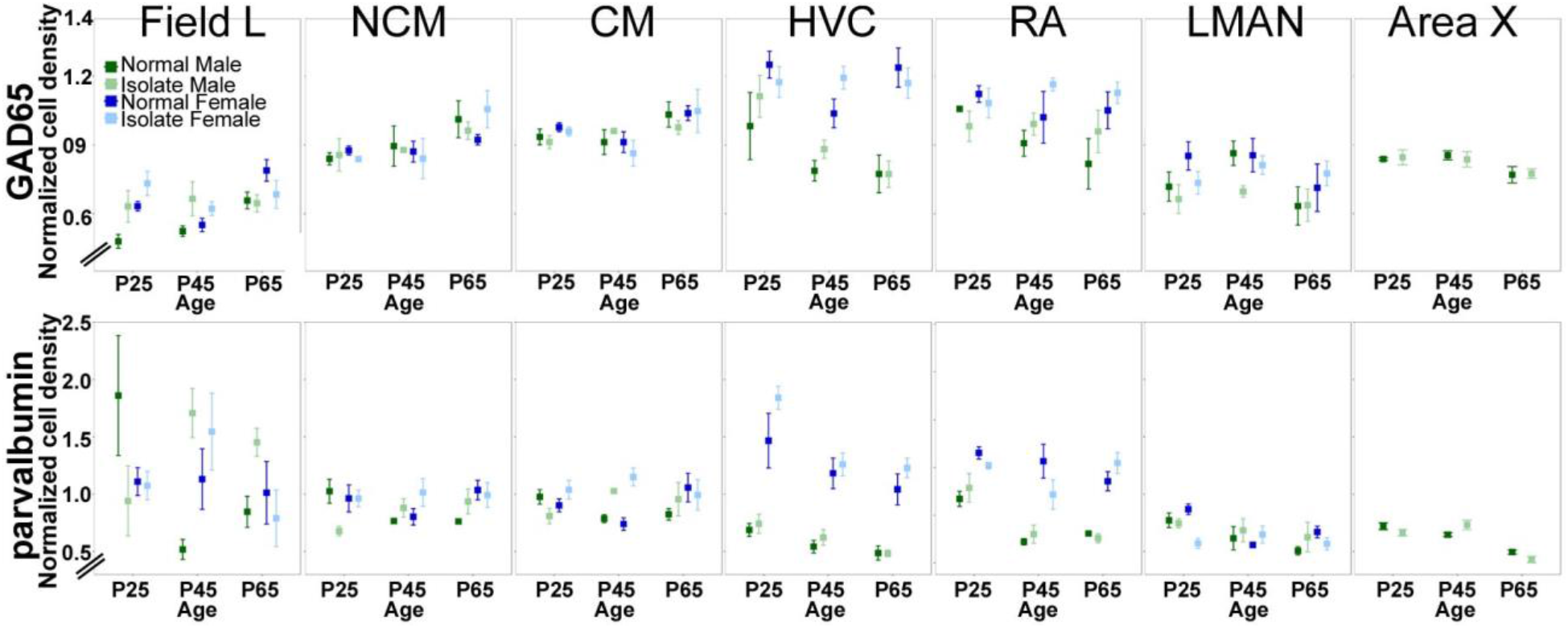
Quantification of inhibitory cell densities across sex, age, tutor experience conditions, and brain area. Quantification of *in situ* hybridization labeling for GAD65 (*top*) and parvalbumin (*bottom*) revealed that cell type densities are determined by age, sex, and tutor experience differently across major nodes of the song network. Plotted are means ± s.e.m.

### Statistics

Statistical tests within each region for main effects of age, sex, condition, and the age * condition interaction were run with the anova function with Type III analysis, in R (R version 3.5.1, stats package v 3.6.0) with α = 0.05. Parametric post hoc analyses were performed using the TukeyHSD function in the stats package.

### Gene expression mapping

To create a comprehensive profile of hybridization across the brain, relative abundance of staining was subjectively determined and divided into four categories: high (+++), medium (++), low (+), and absent (−−). First, brain areas of consistently high and low staining were identified to serve as standards for each gene. Then, a brain region that showed an intermediate hybridization level was designated medium. These standards held for one gene across age, sex, and rearing environment but were not necessarily the same between GAD65 and parvalbumin datasets. Observations across multiple sections were averaged within an individual. Hybridization staining was mapped across the entire brain for subjective and relative intensity of staining using published and online canary and zebra finch atlases as guides, and the revised avian brain nomenclature (the ZEBrA database, Oregon Health & Science University, Portland, OR 97239; http://www.zebrafinchatlas.org; Stokes et al., 1974; Reiner et al., 2004; Nixdorf-Bergweiler and Bischof, 2007)

## Results

All main effect and interaction statistical results are listed in **Table 1**, and guides to quantified neuroanatomical regions are in **Fig 3**.

**Table 1.**
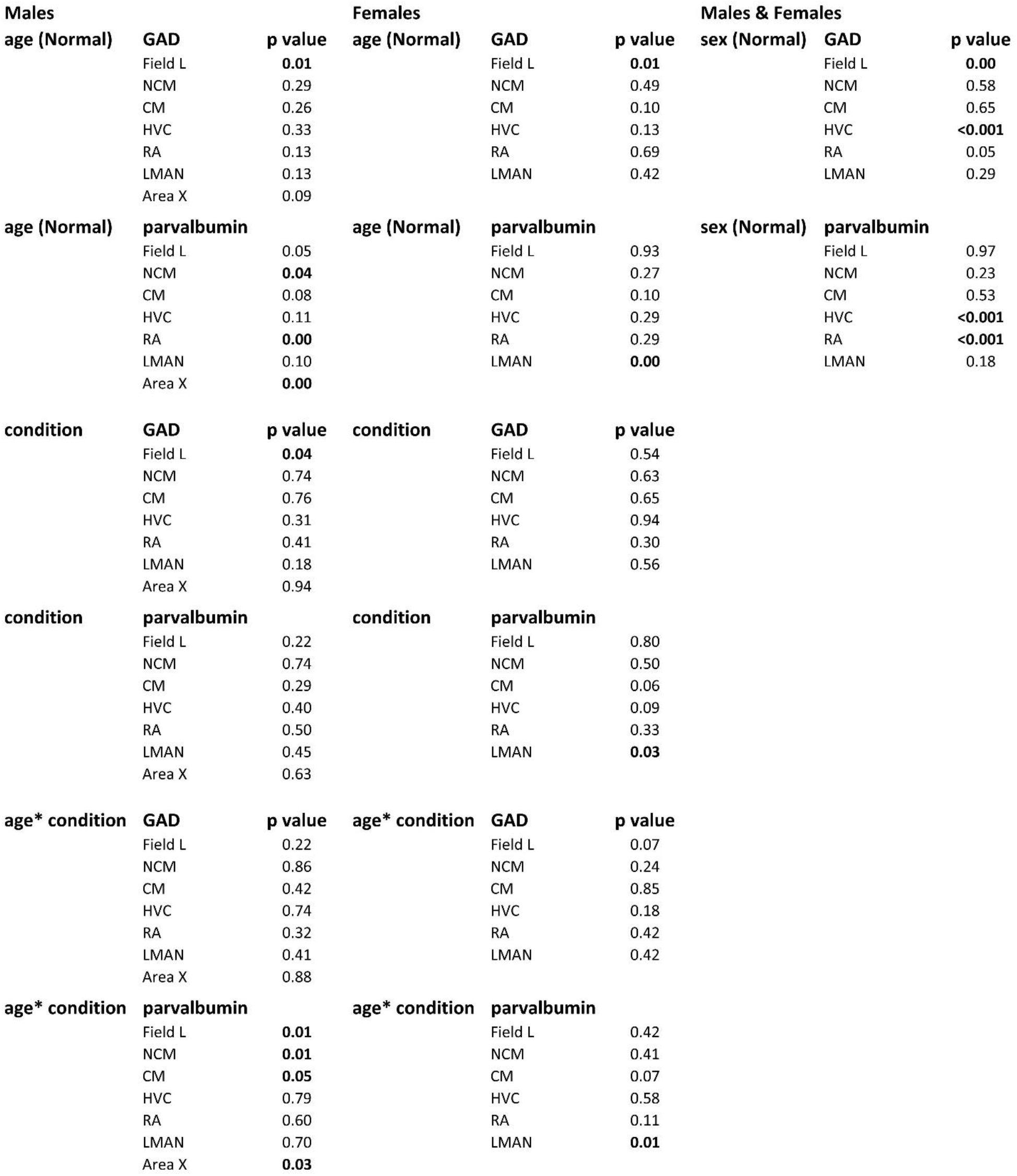
Statistical results. p values for all main effect comparisons and interaction tests between age and tutor experience. Age comparisons were made only for Normal males and females. “Condition” is the comparison of Normal and Isolate birds.

| Males |  |  | Females |  |  | Males & Females |  |  |
| --- | --- | --- | --- | --- | --- | --- | --- | --- |
| age (Normal) | GAD | p value | age (Normal) | GAD | p value | sex (Normal) | GAD | p value |
|  | Field L | **0.01** |  | Field L | **0.01** |  | Field L | **0.00** |
|  | NCM | 0.29 |  | NCM | 0.49 |  | NCM | 0.58 |
|  | CM | 0.26 |  | CM | 0.10 |  | CM | 0.65 |
|  | HVC | 0.33 |  | HVC | 0.13 |  | HVC | **<0.001** |
|  | RA | 0.13 |  | RA | 0.69 |  | RA | 0.05 |
|  | LMAN | 0.13 |  | LMAN | 0.42 |  | LMAN | 0.29 |
|  | Area X | 0.09 |  |  |  |  |  |  |
| age (Normal) | parvalbumin |  | age (Normal) | parvalbumin |  | sex (Normal) | parvalbumin |  |
|  | Field L | 0.05 |  | Field L | 0.93 |  | Field L | 0.97 |
|  | NCM | **0.04** |  | NCM | 0.27 |  | NCM | 0.23 |
|  | CM | 0.08 |  | CM | 0.10 |  | CM | 0.53 |
|  | HVC | 0.11 |  | HVC | 0.29 |  | HVC | **<0.001** |
|  | RA | **0.00** |  | RA | 0.29 |  | RA | **<0.001** |
|  | LMAN | 0.10 |  | LMAN | **0.00** |  | LMAN | 0.18 |
|  | Area X | **0.00** |  |  |  |  |  |  |
| condition | GAD | p value | condition | GAD | p value |  |  |  |
|  | Field L | **0.04** |  | Field L | 0.54 |  |  |  |
|  | NCM | 0.74 |  | NCM | 0.63 |  |  |  |
|  | CM | 0.76 |  | CM | 0.65 |  |  |  |
|  | HVC | 0.31 |  | HVC | 0.94 |  |  |  |
|  | RA | 0.41 |  | RA | 0.30 |  |  |  |
|  | LMAN | 0.18 |  | LMAN | 0.56 |  |  |  |
|  | Area X | 0.94 |  |  |  |  |  |  |
| condition | parvalbumin |  | condition | parvalbumin |  |  |  |  |
|  | Field L | 0.22 |  | Field L | 0.80 |  |  |  |
|  | NCM | 0.74 |  | NCM | 0.50 |  |  |  |
|  | CM | 0.29 |  | CM | 0.06 |  |  |  |
|  | HVC | 0.40 |  | HVC | 0.09 |  |  |  |
|  | RA | 0.50 |  | RA | 0.33 |  |  |  |
|  | LMAN | 0.45 |  | LMAN | **0.03** |  |  |  |
|  | Area X | 0.63 |  |  |  |  |  |  |
| age* condition | GAD | p value | age* condition | GAD | p value |  |  |  |
|  | Field L | 0.22 |  | Field L | 0.07 |  |  |  |
|  | NCM | 0.86 |  | NCM | 0.24 |  |  |  |
|  | CM | 0.42 |  | CM | 0.85 |  |  |  |
|  | HVC | 0.74 |  | HVC | 0.18 |  |  |  |
|  | RA | 0.32 |  | RA | 0.42 |  |  |  |
|  | LMAN | 0.41 |  | LMAN | 0.42 |  |  |  |
|  | Area X | 0.88 |  |  |  |  |  |  |
| age* condition | parvalbumin |  | age* condition | parvalbumin |  |  |  |  |
|  | Field L | **0.01** |  | Field L | 0.42 |  |  |  |
|  | NCM | **0.01** |  | NCM | 0.41 |  |  |  |
|  | CM | **0.05** |  | CM | 0.07 |  |  |  |
|  | HVC | 0.79 |  | HVC | 0.58 |  |  |  |
|  | RA | 0.60 |  | RA | 0.11 |  |  |  |
|  | LMAN | 0.70 |  | LMAN | **0.01** |  |  |  |
|  | Area X | **0.03** |  |  |  |  |  |  |

**Figure 3.**
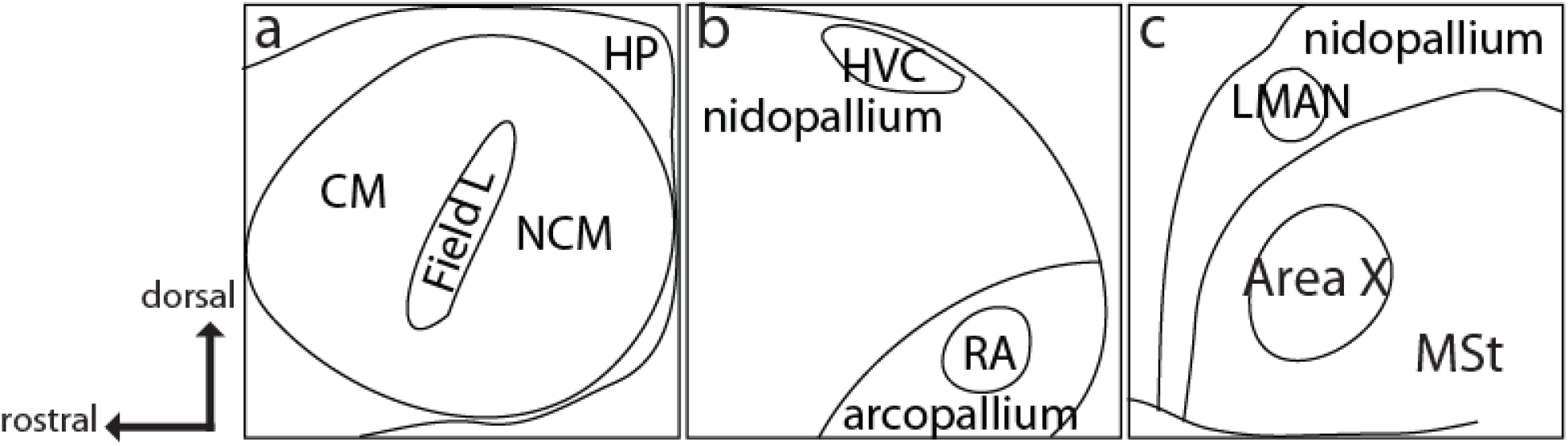
Schematics of song network areas and adjacent areas used for quantification, depicted as in representative *in situ* images. **a)** auditory forebrain showing three major components Field L, NCM, CM, and adjacent hippocampus (HP). **b)** telencephalic region of nidopallium and arcopallium that include HVC and RA, respectively. **c)** LMAN is positioned within the rostral nidopallium, and Area X within the medial striatum (MSt).

### In males, significant effects of age and age * condition interaction are detected only in Field L for GAD65

The only brain region with statistically significant main effect across the three ages in Normal males is Field L (F_(2,6)_ = 9.38, p = 0.01); pairwise posthoc comparisons revealed P65 is significantly different from P25 (p = 0.02) and P45 (p = 0.04) density measures. Similarly, the only brain region with a main effect of tutor experience in males is Field L (F_(1,12)_ = 5.48, p = 0.04; **Fig 2, 4**). There were no significant interactions between age and condition in males, although we observed a trend towards increasing density across age through P65 in Normal males in auditory regions Field L, NCM, CM but decreasing in sensorimotor and motor regions HVC, RA, LMAN, and Area X (**Fig 2**).

**Figure 4.**
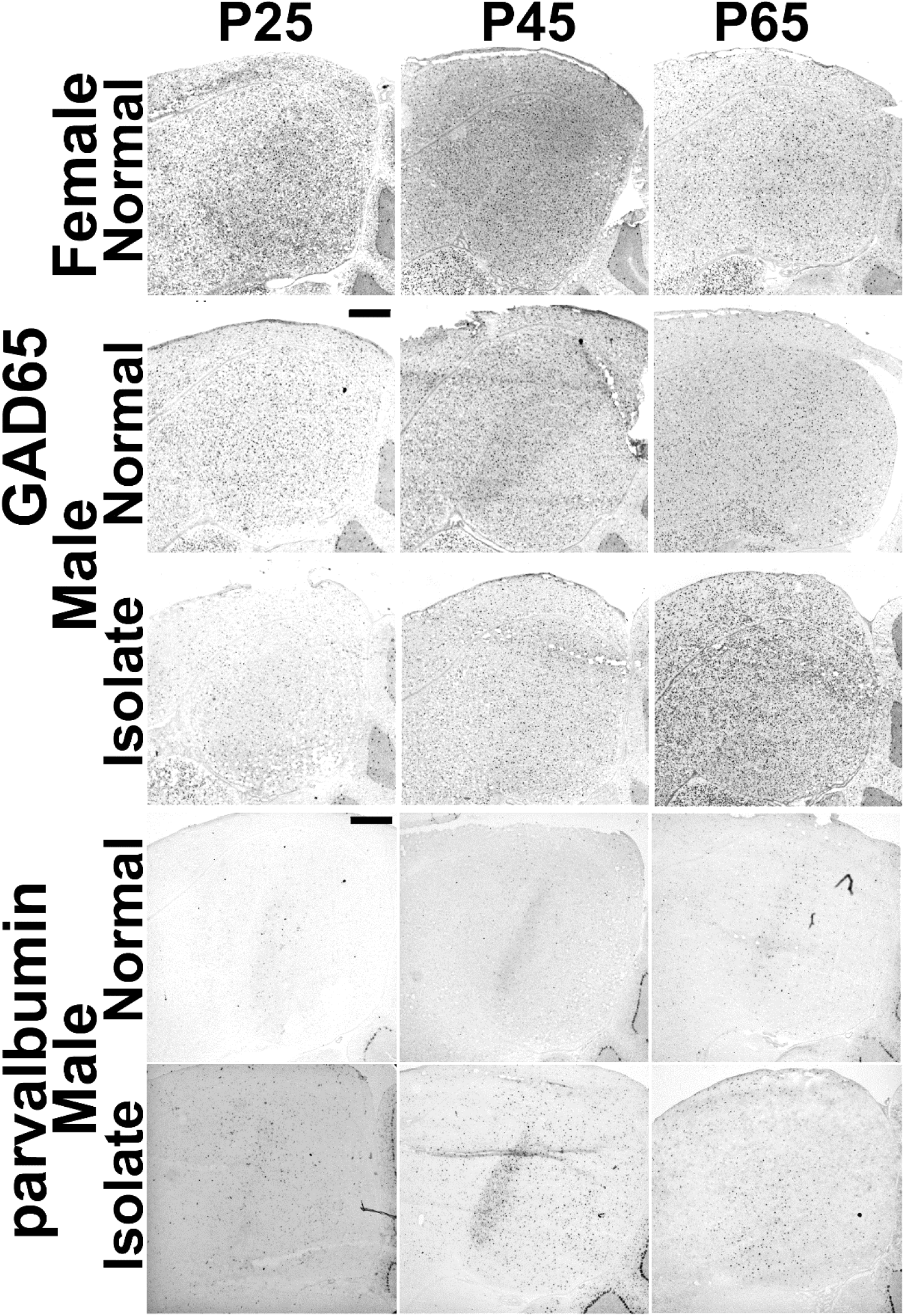
Representative images of auditory forebrain showing GAD65 and parvalbumin. Images show major significant statistical effects between Normal and Isolate males for both genes, and between Normal males and females for GAD65. Scale bars = 500 μm.

### In males, significant effects of age and tutor experience are distributed across the song network for parvalbumin

Several brain areas show significant changes in Normal males across the three ages sampled. In the auditory forebrain, Field L and NCM showed significant age fluctuations (Field L: (F_(2,6)_ = 4.91, p = 0.05, posthoc test showed P25 and P45 are different, p = 0.05; NCM: F_(2,6)_ = 5.79, p = 0.04; posthoc tests revealed marginal differences between P25 and P45 (p = 0.06) and P65 (p = 0.06)), though CM did not reach significance threshold (p = 0.08). In the Anterior Forebrain Pathway, only Area X showed significant differences by age (F_(2,6)_ = 25.86, p = 0.001), with P65 densities distinct from those at P25 (p = 0.001) and P45 (p = 0.01). Parvalbumin cell densities in RA also showed an effect of age (F_(2,6)_ = 22.42, p = 0.002), with densities higher at P25 than the older ages (P25-P45: p = 0.002; P25-P65: p = 0.005; **Fig 2**). There were no significant main effects of tutor experience on any brain area (Table 1).

Several brain areas, however, revealed significant interactions between age and condition in males. All three major components of the auditory forebrain had significant interactions (field L: F_(2,12)_ = 7.85, p = 0.007; NCM: F_(2,12)_ = 7.72, p = 0.007; CM: F_(2,12)_ = 3.94, p = 0.05. In all three components, Normal densities are higher than Isolate densities at P25, but lower than Isolate densities at P45 and P65 (**Fig 2, 4**). Area X also had a significant interaction (F_(2,12)_ = 4.50, p = 0.03); Normal densities were greater than Isolate densities at P25 and P65, but lower than Isolate densities at P45 (**Fig 2**).

### Sex significantly affects cell densities in motor pathway nodes and Field L

We then asked how GAD65 and parvalbumin cell densities compared between males and females, because males but not females sing and it is an open question about how many and which differences in the song network contribute to the behavioral sex difference (Ahmadiantehrani and London, 2017; London, 2017; Shaughnessy et al., 2019).

When we compared Normal males and females, we found significant main effects of sex in Field L (F_(1,12)_ = 15.50, p = 0.002), HVC (F_(1,12)_ = 21.48, p < 0.001), and RA (F_(1,12)_ = 4.73, p = 0.05032) for GAD65 cell densities, and in HVC (F_(1,12)_ = 38.05, p = 4.812e-05) and RA (F_(1,12)_ = 65.78, p = 3.267e-06) for parvalbumin cell densities. In all cases, females had greater cell densities than males (**Fig 2, 5**).

**Figure 5.**
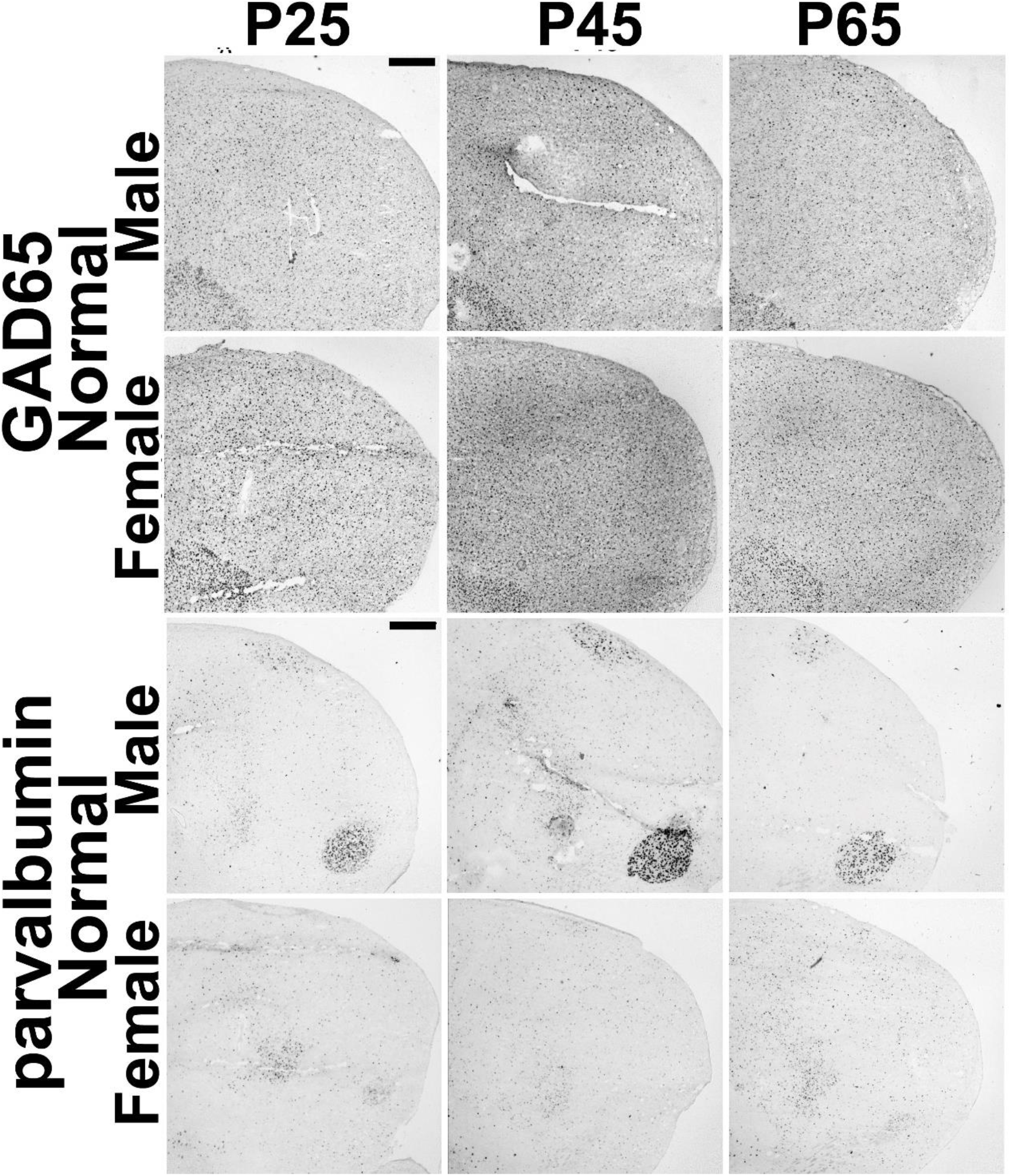
Representative images of HVC and RA showing GAD65 and parvalbumin. Images show major significant statistical effects between Normal males and females and across age. Scale bars = 500 μm.

We also tested for an interaction of age and sex between Normal males and females, and found a significant effect in GAD65 densities only in Field L (F_(4,12)_ = 11.11, p < 0.001), and significant effects on parvalbumin densities in Field L (F_(4,12)_ = 3.23, p = 0.05), CM (F_(4,12)_ = 3.56, p = 0.04), RA (F_(4,12)_ = 4.51, p = 0.02), and LMAN (F_(4,12)_ = 6.19, p = 0.006), with a trend in NCM (F_(4,12)_ = 2.91, p = 0.07) but not HVC (F_(4,12)_ = 1.69, p = 0.22; **Fig 2**).

### Cell densities in female song network areas are minimally affected by age and tutor experience

Finally, we asked if age and tutor experience affected the density of GAD65 and parvalbumin-expressing cells in females. When we compared Normal females by age, we found two significant effects, one for GAD65 (Field L: F_(1,12)_ = 12.46, p = 0.007, posthoc tests showed P65 densities were significantly different from those at P25 (p = 0.04) and P45 (p = 0.006) measures) and one for parvalbumin (LMAN: F_(4,12)_ = 14.94, p = 0.005, posthoc tests showed P25 densities significantly different from those at P45 (p = 0.004) and at P65 (p = 0.03)). The Normal and Isolate females had significantly different parvalbumin-expressing cell densities in LMAN only (F_(1,12)_ = 6.24, p = 0.03; **Fig 2, 6**). The female age * condition interaction was also significant for parvalbumin-expressing cells (F_(2,12)_ = 7.33, p = 0.008; **Fig 2**).

**Figure 6.**
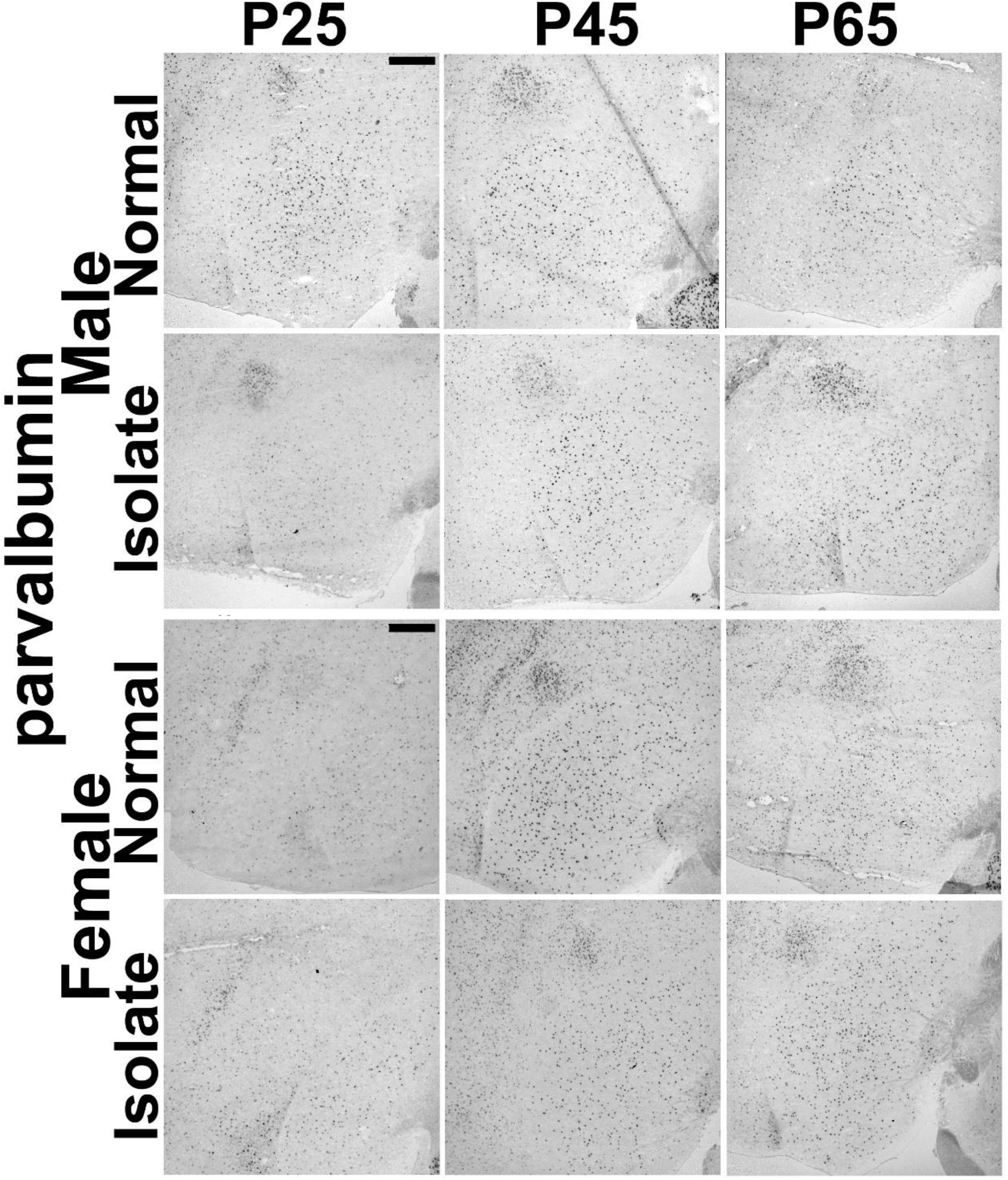
Representative images of parvalbumin-labeled cells in male Area X and female LMAN. Images show comparison between Normal and Isolate birds across age. Scale bars = 500 μm.

### Neuroanatomical distribution and subjective quantification across whole brains

Subjective mappings of *in situ* hybridization labeling revealed widespread distribution of GAD65- and parvalbumin-expressing cells throughout the brain. As a resource, comprehensive tables for both genes, in Normal and Isolate males and females, across all three ages, are included as Supplementary Materials.

## Discussion

Effective behavioral acquisition requires that brain areas are organized appropriately to process information effectively. E/I balance underlies much of this progression, especially during development, because it facilitates specificity in response magnitudes and selectivity. Inhibitory cell types, GAD65 and parvalbumin in particular, are sufficient to gate an extreme example of developmental switches in experience-dependent plasticity: CP. Here, we asked if the density of these two cell types in major areas of a neural network required for developmental song learning in the zebra finch songbird are consistent with a CP for tutor song memorization and the distributed learning that occurs throughout the network.

Our primary prediction was that higher order auditory association areas required for tutor song memorization, NCM and CM, would show age and experience-dependent changes in males. Our data do not fully support this prediction. We found no significant differences based on age or tutor experience condition for GAD65-expressing cells. We did detect a significant effect of age in NCM and a trend in CM for parvalbumin-expressing cell densities, but no effect of condition. In Field L, however, we found significant effects of age and tutor experience on GAD65-expressing cells and of age for parvalbumin-expressing cells. We also found a significant interaction between age and tutor experience condition in male Field L; densities at P25 are higher in Normal compared to Isolate males, but this relationship is flipped at P45 and P65.

The differences found in Field L raise the possibility that primary sensory cortices may be particularly sensitive to the effects of age and experience; the bulk of the work describing functionality of GAD65 and parvalbumin cell subtypes in CPs comes from the primary visual cortex in mammals (Wiesel and Hubel, 1963; Blakemore and Cooper, 1970; Takesian and Hensch, 2013) where GAD65 signaling and parvalbumin cell populations have direct influence on neural plasticity (Hensch et al., 1998; Davis et al., 2015). The current results are intriguing because on the one hand, Field L does not demonstrate distinct response magnitude or selectivity properties for tutor song compared to conspecific song in normally-reared males, yet early song experience does alter the balance of responsiveness and selectivity to song compared to non-song sounds in Field L (Gehr et al., 2000; Maul et al., 2010; Woolley et al., 2010). Notably, other studies that measured parvalbumin protein-labeled cells did not detect alterations in Normally reared males across age (Braun et al., 1991; Cornez et al., 2017). It can, however, be difficult to link different levels of analysis with simple linear relationships. For example, hearing song evokes the same electrophysiological responsiveness in auditory forebrain neurons at P20 and P30, but results in a significant difference in molecular cascade responses between P23 and P30 (Stripling et al., 2001; Ahmadiantehrani and London, 2017). Additional work to link cell populations with activity metrics is needed to reconcile these apparent discrepancies. However, we also note that Field L is highly interconnected with NCM and CM, and may thus still exert an influence on higher order sensory integrating processing (Theunissen et al., 2004).

Interestingly, in Normal males the qualitative trend for GAD65-expressing cells in the three major regions of the auditory forebrain was for density to increase from P25 to P65, but parvalbumin-expressing cell densities were highest at P25 compared to P45 and P65. Perhaps this reflects shifting proportions of inhibitory cell subtypes. In addition to parvalbumin, evidence for somatostatin and VIP inhibitory cell subtype involvement in CP plasticity is accumulating (Trachtenberg, 2015; van Versendaal and Levelt, 2016; Hensch and Quinlan, 2018). Because we performed *in situ* hybridization for both GAD65 and parvalbumin on the same brains, on adjacent sections, these patterns accurately reflect relative population changes within individuals. Assays for other inhibitory cell subtypes would test for the possibility that, for example, somatostatin or VIP cell populations were increasing as parvalbumin subtypes decline.

We were interested in inhibitory cell populations in the female auditory forebrain because it is unclear whether or not females, like males, can memorize tutor song but simply can’t “tell” us because they can’t sing a representation of this memory. Alternatively, tutor song memorization might be male-specific, a reflection of a set of cell types or experience-dependent activation profiles in the auditory forebrain that are sexually dimorphic. Juvenile females can discriminate songs and maintain a trace of their dad’s songs into adulthood (Braaten et al., 2006; Riebel, 2009), but interestingly, while early song isolation creates deficits in behavioral discrimination of songs, distinct electrophysiological responses in Field L to hearing song only occurred in song-isolated females and not in control females (Lauay et al., 2004; Woolley et al., 2010; Diez et al., 2017). The bulk of molecular evidence is that there is are no sex differences in molecular responses to hearing song (but see Bailey and Wade, 2003), but recent data do reveal a significant sex difference at P30 but not before, coincident with the onset of the CP for tutor song memorization (Ahmadiantehrani and London, 2017).

Here, we did not detect a significant effect of condition in any auditory forebrain component in females. In Normal females, the only significant effect of age was for GAD65-expressing cells in Field L. Females, like males, had higher cell densities at P65 compared to P25 and P45; there was also a significant sex difference in GAD65-expressing cells in Field L. This pattern may thus represent an inhibitory cell population with different overall levels in males and females, but which is regulated similarly by experience-independent maturational mechanisms.

Although the study design was focused on the CP for tutor song memorization, the auditory forebrain projects to nodes in the Anterior Forebrain and Posterior Motor Pathways (Vates et al., 1996; Bauer et al., 2008; Mandelblat-Cerf et al., 2014). It was therefore possible that song isolation, by preventing the sensorimotor error correction process, for example, would affect inhibitory cell populations throughout the network.

While we did find significant effects of age in Normal males, with higher parvalbumin-expressing cell densities at P25 than at P45 and P65 in RA, and a denser population at P65 compared to P25 and P45 in Area X, we found no main effect of tutor experience on any component of the Anterior Forebrain or Posterior Motor Pathways in males. This finding is consistent with prior work showed almost no genomic effect of song isolation in these areas (Mori and Wada, 2015). It is, however, possible that our ages were too early to see the effects of song isolation on inhibitory cell populations in the sensorimotor and motor network. For example, in HVC, densities of parvalbumin cells are higher in Isolate males than Normal males by P120, but not at P60 in HVC (Cummings, 2016).

Both HVC and RA had significant sex differences in GAD65- and parvalbumin-expressing cells. In all cases, females had higher densities than males. Lower levels of inhibitory signaling are more permissive for neural plasticity in HVC (Vallentin et al., 2016). If indeed sparser inhibitory cell population is permissive for flexible syllables, then perhaps the greater density in females is part of the cellular substrate that does not allow learned song production at all in females.

Within females, we did find main effects of age and condition, and in their interaction, on parvalbumin-expressing cell densities within the Anterior Forebrain Pathway node LMAN. We found no such effects in males. The effects derive primarily from differences at P25. It is possible that one of the underlying mechanisms that alters the volume of LMAN from P25 to P45 is a decrease in the inhibitory cell population (Bottjer et al., 1985; Bottjer and Sengelaub, 1989; Korsia and Bottjer, 1989; Johnson and Bottjer, 1994).

In fact, we observed that several measures showed apparent magnitude differences between Isolate and Normal birds at P25. This is interesting because we removed birds from their home aviaries at P21 to create the Isolate group, whereas the Normal birds remained undisturbed in their home aviaries. While formally possible that a dramatic change in cell subtype populations could occur based on differential song exposure within four days, it is also possible that the distinctions at P25 is a result of this social stressor. Future experiments could employ more balanced control conditions successfully used to demonstrate epigenetic influences of tutor experience: Tutored condition with one adult male and one adult female in comparison to the two adult female Isolate condition, as used here. This comparison thus controls for aviary removal and standardizes social complexity while still varying tutor experience. There is precedence for these conditions causing distinctions in epigenetic- and systems-levels of analysis in juveniles males (Kelly et al., 2018; Layden et al., 2019).

Our initial predictions were based on the evidence that the NCM and CM portions of the auditory forebrain are required for tutor song memorization and that this is the type of learning that is gated by a CP (Phan et al., 2006; London and Clayton, 2008; Gobes et al., 2010; Yanagihara and Yazaki-Sugiyama, 2016; Ahmadiantehrani and London, 2017; London, 2017). But largely, it was Field L, not NCM and CM, that showed effects of age and tutor experience. Perhaps our predictions relied too strictly on CP mechanisms described for a CP in visual field organization, which is distinct from tutor song memorization in several ways.

First, the experiential conditions that gate CP plasticity is sensory deprivation in the primary visual cortex but the CP for tutor song memorization is closed only by exposure to a tutor; female calls, which have very similar acoustic features to simple song elements, and the bird’s own song, are not sufficient auditory stimuli to close the CP (Eales, 1985, 1987; Slater et al., 1988; Morrison and Nottebohm, 1993; Hensch, 2005). Second, the CP for tutor song memorization is for learning, not perception. In fact, tutor song memorization is not rote, it is optimal when song is presented in social interactions and multimodal sensory experiences (Adret, 2004; Deregnaucourt et al., 2013). Third, a feature of the CP in primary visual cortex is that extended experience-dependent plasticity is a delay of maturation, but epigenetic evidence from juvenile male auditory forebrain is not consistent with this idea (Takesian and Hensch, 2013; Kelly et al., 2018). Fourth, the cellular organization of avian brains is not the same as in mammals. Much of the study of local inhibitory circuits in mammals is related to their effect on pyramidal cells. While gene analysis has uncovered strong parallels between bird forebrain and mammalian neocortex, cell types are organized distinctly, and more distributed in bird brains (Dugas-Ford et al., 2012; Karten, 2013; Calabrese and Woolley, 2015). Thus local circuits may also require a different configuration to have similar effects. On the other hand, both Field L and primary visual cortex in mammals receive major thalamic inputs, which are essential for patterning the receptive fields of sensory cells. It is possible that this property explains the effects we detected here (Theunissen et al., 2004).

The current study represents an initial foray into assessing the potential role of local inhibitory circuits in the emergent response and selectivity properties across the brain network required for developmental song learning in zebra finches. The results indicate specific alterations to the density of inhibitory cell subtypes depending on the age, sex, and tutor experience of the individual, and differed based on the brain region quantified. Important next steps include investigation of additional cell subtypes and the measurement of dynamic synaptic function of these networks. The zebra finch song network has already provided key insights into neural properties that promote and limit the ability to learn and produce complex behaviors; additional investigation into the role of local inhibitory circuits is thus likely to be fruitful.

## Conflicts of interest

The authors declare no conflicts of interest.

## Acknowledgments

We thank Sarah Greta for assistance with image quantification. Supported by the University of Chicago and the Institute for Mind and Biology.

