## Supplemental Tables for "Inhibitory cell populations depend on age, sex, and prior experience across a neural network for Critical Period learning"

### Supplementary tables

Tables of qualitative brain-wide expression mapping.

**Table** **page**

Abbreviations for neuroanatomical nomenclature used in subsequent tables ………………. 2

Whole brain GAD65 mapping ………………………………………………………………..………………. 3

Song network GAD65 mapping ………………………..…………………………………..………………. 4

Whole brain parvalbumin mapping ………….………………………………………………..……………. 5

Song network parvalbumin mapping ……………..……………………………………..………………. 6

**Telencephalon (pallium, subpallium)**

| | |
|---|---|
| HA | Apical hyperpallium |
| HD | Dorsal hyperpallium |
| M | Mesopallium |
| N | Nidopallium |
| S | Septum |
| A | Acropallium |
| E | Ectopallium |
| LSt (PA) | Lateral striatum |
| GP (PP) | Globus pallidus |
| SL | Lateral septal nucleus |
| SM | Medial septal nucleus |
| MSt | Medial striatum |
| Tn | Nucleus taeniae |
| B | Nucleus basalis |
| AL | Auditory lobule |
| LMAN | Magnocellular nucleus of the anterior nidopallium |
| Area X | Proper name |
| RA | Robust nucleus of the arcopallium |
| HVC | Proper name |
| Field L | Proper name |
| Bas | Basorostral nucleus |
| OM | Occipitomesencephalic tract |
| CDLco | Core of the dorsolateral corticoid area |
| Alat | Lateral acropallium |

**Diencephalon (thalamus/hypothalamus)**

| | |
|---|---|
| PVM (PVN) | Paraventricular nucleus |
| LHy | Lateral hypothalamus |
| GLv | Ventral part of the lateral geniculate nucleus |
| Rot/Rt | Nucleus rotundus |
| DLA | Anterior nucleus of the dorsolateral thalamus |
| Ov/ovco | Core of the nucleus ovoidalis |
| DLMco | Core of the medial part of the dorsolateral nucleus of the anterior thalamus |
| DP/DLP | Dorsolateral nucleus in the posterior thalamus |
| SpM | Medial spiriform |
| SpL | Lateral spiriform |
| EM | Ectomamillary nucleus |
| DMP | Posterior dorsomedial nucleus |
| HM/HL | Medial/lateral habenula |
| SPC | Parvocellular part of the superficial nucleus of the thalamus |
| SP/SPT | Nucleus subpretectalis |
| Pt | Pretectal nucleus |
| SubG | Subgeniculate nucleus |
| ALP | Posterior nucleus of ansa lenticularis |
| TSM | Septomesencephalic tract |
| OvPG | Oval pregeniculate nucleus |
| Ret | Reticular nucleus of the thalamus |
| SRt | Nucleus subrotundus |
| EPD | Dorsal part of the entopeduncular nucleus |
| VMH | Ventromedial nucleus of the hypothalamus |
| DMA | Dorsomedial nucleus of the anterior thalamus |
| DLM | dorsolateral thalamus |

**Mesencephalon (midbrain)**

| | |
|---|---|
| LM | Lentiform nucleus |
| VTA (AVT) | Ventral tegmetal area |
| Ico/MLd | Intercollicular nucleus/dorsal part of the lateral mesencephalic nucleus |
| IM/IPC | Magnocellular isthmus nuclei/parvocellular isthmus nuclei |
| SGC | Stratum griseum centrale |
| SGP/SGF | Substantia grisea et fibrosa periventricularis |
| LoC | Locus coeruleus |
| EW | Nucleus of Edinger-Westphal |
| Ru | Red nucleus |
| NIII | Nucleus of III nerve |
| PL/PM | Lateral/medial pons |
| nIV | Nucleus of IV nerve |
| FLM | Medial longitudinal fascicle |
| SN | Substantia negra |
| LLD | Dorsal part of the lateral lemniscal nucleus |
| ISo | Isthmooptic nucleus |
| PAG | Periaqueductal gray |

**Metencephalon (pons, cerebellum)**

| | |
|---|---|
| LA | Nucleus laminaris |
| FLM | Medial longitudinal fascicle |
| RPgc | Reticular nucleus of caudal pons, gigantocellularis |
| Dcn | Deep cerebellar nuclei |
| Vest | Vestibular nuclei |
| MC | Magnocellular nucleus |
| Ang | Angular nucleus |
| nV | Principle sensory nucleus of the trigeminal nerve |
| n7 | Nucleus of the facial nerve |

**Myelencephalon (medulla)**

| | |
|---|---|
| Vest | Vestibular nuclei |

GAD65

| Brain Area | P25 | | | | P45 | | | | P65 | | | |
|---|---|---|---|---|---|---|---|---|---|---|---|---|
| | Male | | Female | | Male | | Female | | Male | | Female | |
| | Normal | Isolate | Normal | Isolate | Normal | Isolate | Normal | Isolate | Normal | Isolate | Normal | Isolate |
| HA | +(++) | +(++) | +(++) | +(++) | +(++) | +(++) | +(++) | +(++) | +(++) | +(++) | +(++) | +(++) |
| M | +(++) | +(++) | +(++) | +(++) | +(++) | +(++) | +(++) | +(++) | +(++) | +(++) | +(++) | +(++) |
| HD | +(++) | +(++) | +(++) | +(++) | +(++) | +(++) | +(++) | +(++) | +(++) | +(++) | +(++) | +(++) |
| N | +(++) | +(++) | +(++) | +(++) | +(++) | +(++) | +(++) | +(++) | +(++) | +(++) | +(++) | +(++) |
| GP (PP) | +++ | ++ | ++ | ++ | +++ | ++ | ++ | +++ | +++ | +++ | ++ | +++ |
| LSt (PA) | ++ | ++ | +(++) | +(++) | +(++) | ++ | +(++) | ++ | ++ | ++ | +(++) | +(++) |
| MSt | +(++) | +(++) | +(++) | +(++) | +(++) | +(++) | +(++) | +(++) | +(++) | +(++) | +(++) | +(++) |
| SL | + | + | + | + | + | + | + | + | + | + | + | + |
| SM | + | + | + | + | + | + | + | + | + | + | + | + |
| A | +(++) | +(++) | +(++) | +(++) | +(++) | +(++) | +(++) | +(++) | +(++) | +(++) | +(++) | +(++) |
| Tn | +(++) | +(++) | +(++) | +(++) | +(++) | +(++) | +(++) | +(++) | +(++) | +(++) | +(++) | +(++) |
| E | ++ | ++ | ++ | ++ | ++ | ++ | ++ | ++ | ++ | + | ++ | ++ |
| B | + | + | + | + | + | + | + | + | + | + | + | + |
| PVM (PVN) | + | + | + | + | + | + | + | + | + | ++ | + | + |
| LHy | ++ | + | ++ | ++ | + | ++ | ++ | + | ++ | ++ | ++ | ++ |
| GLv | +(++) | + | +(++) | + | +(++) | +(++) | +(++) | +(++) | + | + | ++ | +(++) |
| Rot/Rt | -- | -- | -- | -- | -- | -- | -- | -- | -- | -- | -- | -- |
| LA | -- | -- | -- | -- | -- | -- | -- | -- | -- | -- | -- | -- |
| DLA | + | ++ | + | + | + | + | ++ | + | + | + | + | + |
| Ov/ovco | -- | -- | -- | -- | -- | -- | -- | -- | -- | -- | -- | -- |
| DLMco | -- | -- | -- | -- | -- | -- | -- | -- | -- | -- | -- | -- |
| DP/DLP | -- | -- | -- | -- | -- | -- | -- | -- | -- | -- | -- | -- |
| SpM | -- | -- | -- | -- | -- | -- | -- | -- | -- | -- | -- | -- |
| S | + | ++ | ++ | ++ | + | + | + | + | + | ++ | + | + |
| SpL | ++ | ++ | ++ | ++ | ++ | ++ | +++ | ++ | ++ | ++ | ++ | + |
| EM | +++ | + | ++ | + | ++ | ++ | ++ | ++ | ++ | ++ | ++ | ++ |
| DMP | -- | -- | -- | -- | -- | -- | -- | -- | -- | -- | -- | -- |
| DLM | -- | -- | -- | -- | -- | -- | -- | -- | -- | -- | -- | -- |
| LM | ++ | + | + | + | + | + | ++ | ++ | ++ | ++ | + | + |
| VTA (AVT) | + | + | ++ | + | + | + | + | + | + | ++ | + | + |
| Ico/MLd | +++ | ++ | +++ | +++ | +++ | +++ | +++ | +++ | +++ | +++ | ++ | +++ |
| IM/IPC | +++ | +++ | +++ | +++ | +++ | +++ | +++ | +++ | +++ | +++ | +++ | +++ |
| HM/HL | -- | -- | -- | -- | -- | -- | -- | -- | -- | -- | -- | -- |
| SGC | + | + | + | + | + | + | + | + | ++ | + | + | + |
| SGP/SGF | ++ | + | ++ | + | ++ | ++ | + | + | ++ | ++ | + | + |
| SPC | -- | -- | -- | -- | -- | -- | -- | -- | -- | -- | -- | -- |
| SP/SPT | +++ | ++ | +++ | ++ | +++ | ++ | ++ | ++ | ++ | ++ | +++ | ++ |
| Pt | ++ | ++ | ++ | ++ | ++ | +++ | +++ | +++ | +++ | +++ | ++ | ++ |
| EW | ++ | ++ | ++ | ++ | ++ | ++ | ++ | ++ | ++ | ++ | + | ++ |
| Ru | + | + | + | + | + | + | + | + | + | + | + | + |
| NIII | -- | -- | -- | -- | -- | -- | -- | + | -- | + | + | -- |
| PL/PM | -- | + | -- | -- | -- | -- | + | + | -- | + | -- | + |
| nIV | -- | -- | -- | -- | -- | -- | -- | -- | -- | -- | -- | -- |
| FLM | -- | -- | -- | -- | -- | -- | -- | -- | -- | -- | -- | -- |
| LoC | ++ | ++ | ++ | ++ | ++ | ++ | + | ++ | + | ++ | ++ | + |
| RPgc | + | + | + | + | + | + | + | + | ++ | + | + | + |
| dcn | ++ | + | +(++) | + | + | + | + | +(++) | +(++) | +(++) | + | +(++) |
| vest | ++ | ++ | ++ | ++ | ++ | ++ | ++ | ++ | + | ++ | ++ | ++ |
| AL | +(++) | +(++) | +(++) | +(++) | +(++) | +(++) | +(++) | +(++) | +(++) | +(++) | +(++) | +(++) |
| LMAN | +(++) | +(++) | +(++) | +(++) | +(++) | +(++) | +(++) | +(++) | +(++) | +(++) | +(++) | +(++) |
| Area X | +(++) | +(++) | x | x | +(++) | +(++) | x | x | +(++) | +(++) | x | x |
| RA | +(++) | +(++) | +(++) | +(++) | +(++) | +(++) | +(++) | +(++) | +(++) | +(+++) | +(++) | +(++) |
| HVC | +(++) | +(++) | +(++) | +(++) | +(+++) | +(+++) | +(++) | +(++) | +(+++) | +(+++) | +(++) | +(++) |
| Field L | +(++) | +(++) | +(++) | +(++) | +(++) | +(++) | ++ | +(++) | + | +(+++) | +(++) | ++ |
| Bas | ++ | ++ | ++ | ++ | +++ | ++ | ++ | ++ | ++ | ++ | ++ | ++ |
| SN | ++ | ++ | ++ | +++ | +++ | ++ | ++ | ++ | ++ | ++ | ++ | ++ |
| MC | -- | -- | -- | -- | + | + | -- | -- | -- | -- | -- | -- |
| SubG | ++ | ++ | ++ | + | ++ | ++ | ++ | ++ | ++ | ++ | ++ | ++ |
| Ang | -- | -- | -- | -- | -- | -- | -- | -- | -- | -- | -- | -- |
| nV | -- | + | -- | -- | -- | -- | -- | -- | -- | -- | -- | -- |
| ALP | +++ | ++ | +++ | +++ | +++ | ++ | +++ | +++ | +++ | +++ | +++ | +++ |
| LLD | ++ | ++ | ++ | + | ++ | ++ | + | + | + | + | ++ | ++ |
| TSM | -- | -- | -- | -- | -- | -- | -- | -- | -- | -- | -- | -- |
| OvPG | ++ | ++ | ++ | ++ | ++ | ++ | +++ | ++ | ++ | ++ | ++ | ++ |
| Ret | +++ | +++ | +++ | +++ | +++ | +++ | +++ | +++ | +++ | +++ | +++ | +++ |
| ISo | -- | -- | -- | -- | -- | -- | -- | -- | -- | -- | -- | -- |
| n7 | + | + | ++ | + | + | + | + | + | + | + | + | + |
| SRt | + | + | ++ | + | + | ++ | ++ | ++ | ++ | + | + | + |
| OM | + | -- | + | x | -- | -- | -- | -- | x | -- | x | -- |
| CDLco | + | + | + | + | + | + | + | + | + | + | + | + |
| EPD | + | + | ++ | + | + | + | + | + | + | + | + | + |
| PAG | ++ | ++ | ++ | ++ | ++ | ++ | ++ | ++ | ++ | ++ | ++ | ++ |
| VMH | + | + | + | + | + | + | + | + | + | + | x | + |
| Alat | +(++) | +(++) | +(++) | + | +(++) | +(++) | +(++) | +(++) | +(++) | +(++) | +(++) | +(++) |

| Key | Standards |
|---|---|
| + = majority of stained cells | + = SM |
| (+) = staining on some cells in area | ++ = LSt (PA) |
| X = area obscured | +++ = Ret |

**GAD65**

| Brain Area | P25 | | | | P45 | | | | P65 | | | |
|---|---|---|---|---|---|---|---|---|---|---|---|---|
| | Male | | Female | | Male | | Female | | Male | | Female | |
| | Normal | Isolate | Normal | Isolate | Normal | Isolate | Normal | Isolate | Normal | Isolate | Normal | Isolate |
| AL | +(++) | +(++) | +(++) | +(++) | +(++) | +(++) | +(++) | +(++) | +(++) | +(++) | +(++) | +(++) |
| LMAN | +(++) | +(++) | +(++) | +(++) | +(++) | +(++) | +(++) | +(++) | +(++) | +(++) | +(++) | +(++) |
| Area X | +(++) | +(++) | x | x | +(++) | +(++) | x | x | +(++) | +(++) | x | x |
| RA | +(++) | +(++) | +(++) | +(++) | +(++) | +(++) | +(++) | +(++) | +(++) | +(+++) | +(++) | +(++) |
| HVC | +(++) | +(++) | +(++) | +(++) | +(+++) | +(+++) | +(++) | +(++) | +(+++) | +(+++) | +(++) | +(++) |
| Area L | +(++) | +(++) | +(++) | +(++) | +(++) | +(++) | ++ | +(++) | + | +(+++) | +(++) | ++ |

| Key | Standards |
|---|---|
| + = majority of stained cells | + = SM |
| (+) = staining on some cells in area | ++ = LSt (PA) |
| X = area obscured | +++ = Ret |

**Parvalbumin**

| Brain Area | P25 | | | | P45 | | | | P65 | | | |
|---|---|---|---|---|---|---|---|---|---|---|---|---|
| | Male | | Female | | Male | | Female | | Male | | Female | |
| | Normal | Isolate | Normal | Isolate | Normal | Isolate | Normal | Isolate | Normal | Isolate | Normal | Isolate |
| HA | + | + | + | + | + | + | + | + | + | + | + | + |
| M | + | + | + | + | + | + | + | + | + | + | + | + |
| HD | + | + | + | + | + | + | + | + | + | + | + | + |
| N | + | + | + | + | + | + | + | + | + | + | + | + |
| GP (PP) | ++ | ++ | ++ | ++ | ++ | ++ | ++ | ++ | ++ | +++ | ++ | ++ |
| LSt (PA) | ++ | ++ | + | ++ | ++ | ++ | ++ | ++ | ++ | ++ | ++ | ++ |
| MSt | + | + | + | + | + | + | + | + | + | + | + | + |
| SL | -- | -- | -- | -- | -- | -- | -- | -- | -- | -- | -- | -- |
| SM | ++ | + | + | + | ++ | + | ++ | + | + | + | + | ++ |
| A | + | + | + | + | + | + | + | + | + | + | + | + |
| Tn | + | + | + | + | + | + | + | + | + | + | + | + |
| E | ++ | ++ | ++ | ++ | ++ | ++ | ++ | ++ | ++ | ++ | ++ | ++ |
| B | -- | -- | -- | -- | -- | -- | -- | -- | -- | -- | -- | -- |
| PVM (PVN) | -- | -- | -- | -- | -- | -- | -- | -- | -- | -- | -- | -- |
| LHy | -- | -- | -- | -- | -- | -- | -- | -- | -- | -- | -- | -- |
| GLv | + | + | + | + | + | + | + | + | + | + | + | + |
| Rot/Rt | +++ | +++ | +++ | +++ | +++ | +++ | +++ | +++ | +++ | ++ | +++ | +++ |
| LA | -- | + | + | -- | + | + | + | + | + | + | -- | + |
| DLA | + | +* | + | + | + | + | + | + | + | + | + | + |
| Ov/ovco | + | + | + | + | + | + | + | + | + | + | + | + |
| DLMco | -- | -- | -- | -- | -- | -- | + | -- | + | + | -- | + |
| DP/DLP | + | + | + | + | ++ | + | + | ++ | ++ | ++ | + | ++ |
| SpM | + | -- | + | -- | ++ | + | -- | + | + | + | + | + |
| SpL | ++ | ++ | +++ | ++ | +++ | +++ | +++ | ++ | ++ | ++ | ++ | ++ |
| EM | + | + | + | + | ++ | ++ | ++ | ++ | ++ | ++ | + | ++ |
| DMP | -- | -- | -- | -- | -- | -- | -- | -- | -- | -- | -- | -- |
| LM | + | + | + | + | + | + | + | + | + | + | + | + |
| VTA (AVT) | -- | -- | -- | -- | -- | -- | -- | -- | -- | -- | -- | -- |
| Ico/MLd | +(++) | +(++) | +(++) | +(++) | +(++) | +(++) | +(++) | +(++) | +(++) | +(++) | +(++) | +(++) |
| IM/IPC | +++ | +++ | +++ | +++ | +++ | +++ | +++ | +++ | +++ | +++ | +++ | +++ |
| HM/HL | -- | -- | -- | -- | + | -- | -- | -- | -- | -- | -- | -- |
| SGC | ++ | ++ | ++ | + | ++ | ++ | + | ++ | ++ | ++ | ++ | + |
| SGP/SGF | ++ | ++ | ++ | + | ++ | ++ | + | ++ | ++ | ++ | ++ | ++ |
| SPC | -- | -- | -- | -- | -- | -- | -- | -- | -- | -- | -- | -- |
| SP/SPT | +++ | ++ | +++ | ++ | +++ | ++ | +++ | ++ | +++ | +++ | +++ | +++ |
| Pt | + | + | + | + | + | + | + | ++ | + | + | + | + |
| EW | -- | -- | -- | -- | + | -- | + | -- | + | + | -- | + |
| Ru | -- | + | + | + | + | + | + | + | + | -- | -- | + |
| NIII | -- | -- | -- | -- | -- | -- | -- | -- | -- | -- | -- | -- |
| PL/PM | + | + | + | + | + | + | -- | + | + | + | + | -- |
| nIV | -- | -- | -- | -- | -- | -- | + | -- | -- | -- | -- | -- |
| FLM | -- | -- | -- | + | -- | -- | -- | -- | -- | -- | -- | -- |
| LoC | + | +* | + | ++ | ++ | ++ | ++ | ++ | ++ | + | + | ++ |
| RPgc | -- | -- | + | + | + | + | -- | + | + | + | -- | + |
| dcn | + | ++ | ++ | + | + | + | + | + | + | + | + | + |
| vest | + | + | + | + | ++ | + | ++ | ++ | ++ | + | + | ++ |
| AL | -- | + | -- | -- | + | -- | + | + | -- | -- | -- | + |
| LMAN | + | + | + | -- | ++ | +(++) | + | ++ | ++(+++) | +(++) | + | + |
| Area X | + | + | X | X | ++ | ++ | X | X | ++ | ++ | X | X |
| RA | + | + | -- | + | +++ | ++ | -- | + | +++ | +++ | -- | -- |
| HVC | +(++) | +(++) | -- | -- | +(+++) | +(+++) | -- | -- | ++(+++) | +(+++) | -- | -- |
| Field L | -- | + | -- | -- | +(++) | -- | + | -- | + | -- | -- | -- |
| Bas | +(++) | +(++) | +(++) | ++ | +(++) | ++ | +(++) | +(++) | +(++) | +(++) | +(++) | +(++) |
| SN | + | ++ | ++ | ++ | +(++) | ++ | ++ | +(++) | ++ | ++ | + | +(++) |
| MC | + | + | ++ | + | ++ | ++ | + | + | ++ | + | + | + |
| SubG | -- | -- | + | -- | -- | -- | -- | -- | -- | -- | -- | -- |
| Ang | ++ | + | ++ | + | ++ | ++ | ++ | ++ | ++ | ++ | ++ | ++ |
| nV | + | + | + | + | + | + | + | + | ++ | + | + | ++ |
| ALP | + | ++ | ++ | + | + | + | ++ | + | ++ | + | + | ++ |
| LLD | + | -- | -- | -- | -- | -- | + | -- | -- | + | -- | ++ |
| TSM | -- | -- | -- | -- | -- | -- | -- | -- | -- | -- | -- | -- |
| OvPG | + | + | + | + | + | + | + | + | + | + | + | + |
| Ret | + | + | + | -- | + | + | -- | + | + | -- | + | + |
| ISo | -- | -- | -- | -- | -- | -- | -- | -- | -- | -- | -- | + |
| n7 | -- | + | + | -- | + | + | + | + | + | + | -- | + |
| SRt | -- | -- | -- | -- | -- | -- | -- | -- | -- | -- | -- | -- |
| DLM | -- | -- | -- | -- | -- | -- | -- | -- | -- | -- | -- | -- |
| CDLco | + | + | + | + | + | + | + | + | + | + | + | + |
| EPD | -- | -- | -- | -- | -- | -- | -- | -- | -- | -- | -- | -- |
| Alat | +(+++) | + | +(++) | +(++) | +(++) | ++(+++) | +(++) | +++ | +(++) | ++(+++) | ++(+++) | +(++) |
| VMH | -- | -- | -- | -- | -- | -- | -- | -- | -- | -- | -- | -- |
| PAG | -- | -- | -- | -- | -- | -- | -- | -- | -- | -- | -- | -- |
| DMA | -- | -- | -- | -- | -- | -- | -- | -- | -- | -- | -- | -- |

| Key | Standards |
|---|---|
| + = staining on majority of cells in area | + = HA |
| (+) = staining on some cells in area | ++ = E |
| X = area obscured | +++ = IM/IPC |

**Parvalbumin**

| Brain Area | P25 | | | | P45 | | | | P65 | | | |
|---|---|---|---|---|---|---|---|---|---|---|---|---|
| | Male | | Female | | Male | | Female | | Male | | Female | |
| | Normal | Isolate | Normal | Isolate | Normal | Isolate | Normal | Isolate | Normal | Isolate | Normal | Isolate |
| **AL** | -- | + | -- | -- | + | -- | + | + | -- | -- | -- | + |
| **LMAN** | + | + | + | -- | ++ | +(++) | + | ++ | ++(+++) | +(++) | + | + |
| **Area X** | + | + | x | x | ++ | ++ | x | x | ++ | ++ | x | x |
| **RA** | + | + | -- | + | +++ | ++ | -- | + | +++ | +++ | -- | -- |
| **HVC** | +(++) | +(++) | -- | -- | +(+++) | +(+++) | -- | -- | ++(+++) | +(+++) | -- | -- |
| **Field L** | -- | + | -- | -- | +(++) | -- | + | -- | + | -- | -- | -- |

| Key | Standards |
|---|---|
| + = staining on majority of cells in area | + = HA |
| (+) = staining on some cells in area | ++ = E |
| X = area obscured | +++ = IM/IPC |
